## Supplementary files for "Micro RNAs are minor constituents of extracellular vesicles and are hardly delivered to target cells"

Manuel Albanese[1,2,3†], Yen-Fu Adam Chen[1,3†], Corinna Hüls[1,3], Kathrin Gärtner[1,3], Takanobu Tagawa[1,3,5], Ernesto Mejias-Perez[2,3], Oliver T. Keppler[2,3], Christine Göbel[1,3], Reinhard Zeidler[1,3,4], Mikhail Shein[6], Anne K. Schütz[6,7] and Wolfgang Hammerschmidt[1,3*]

[1]Research Unit Gene Vectors, Helmholtz Zentrum München, German Research Center for Environmental Health, Munich, Germany

[2]Max von Pettenkofer Institute and Gene Center, Virology, National Reference Center for Retroviruses, Faculty of Medicine, LMU München, Munich, Germany.

[3]German Centre for Infection Research (DZIF), Partner site Munich, Germany

[4]Department of Otorhinolaryngology, Klinikum der Universität München, Marchioninistr. 15, 81377 Munich, Germany

[5]Current address: HIV and AIDS Malignancy Branch, Center for Cancer Research, National Cancer Institute, National Institutes of Health, Bethesda, Maryland, USA

[6] Bavarian NMR Center, Department of Chemistry, Technical University of Munich, 85748, Garching, Germany.

[7]Institute of Structural Biology, Helmholtz Zentrum München, 85764 Neuherberg, Germany.

[†]shared authorship

*Corresponding author

Wolfgang Hammerschmidt

Keywords: extracellular vesicles, herpesvirus, microRNA, fusion assay

**Supplementary Figure S1**

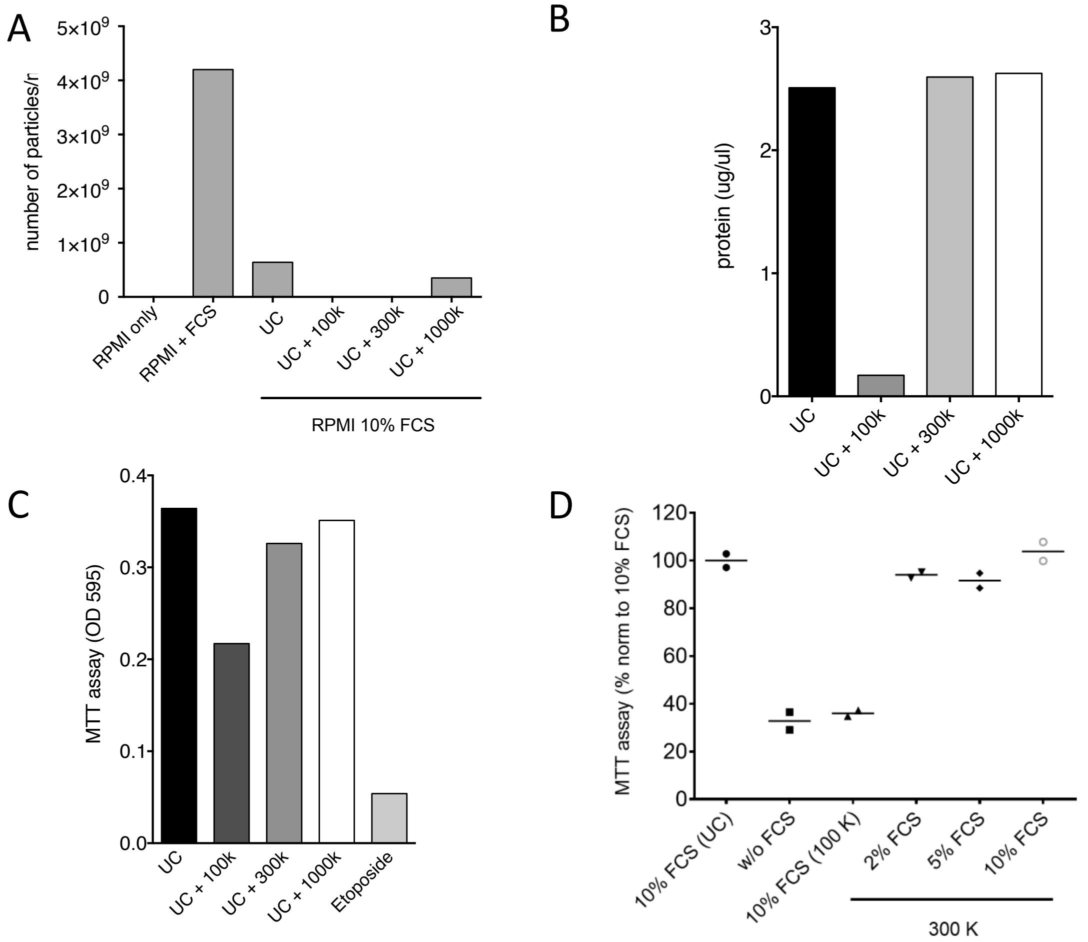


**Preparation of fetal calf serum (FCS) to reduce the concentration of bovine EVs**.

FCS was diluted 1:1 with RPMI1640 medium and centrifuged at 100,000 g at 4 °C in a swinging-bucket rotor (SW28 or 32, Beckman Coulter) for 18 h. After ultracentrifugation (UC), the supernatant was further processed using ultrafiltration spin columns purchased from different manufacturers and with different cutoffs: 100K Amicon Ultra-15 (ultracel regenerated cellulose, Merck), 300K Vivaspin 20 or 1000K Vivaspin 20 concentrators (PES, polyethersulfon, Sartorius). The columns were centrifuged at 2,000 g, 10 °C for 20-30 min. The different preparations were tested for their EV concentrations by NTA, protein content and adverse effects on a lymphoblastoid cell line (LCL). **(A)** EV particle concentrations of RPMI supplemented with 10 % untreated, not EV-depleted fetal calf serum (FCS) or supplemented with 10 % FCS treated with three different centrifugal filter were measured by NTA with the ZetaView instrument PMX110 to analyze the EV concentrations after UC and followed by ultrafiltration as indicated. A single UC step reduced the concentration of EVs to about 20 %, whereas ultrafiltration with 100K or 300K filters removed the majority of bovine EVs. **(B)** Protein concentrations (as measured by Bradford) of the samples analyzed in panel A. Filtration with the 100K filter device led to a considerable reduction of proteins contained in FCS. **(C)** LCLs were cultivated in RPMI1640 supplemented with 10 % FCS conditioned as shown in panel A. After 3 days, an MTT assay was performed to assess the viability of the cells. As a negative control, LCLs were treated with 10 µM etoposide for 1 h to induce cell death. Cells cultivated in RPMI1640 with 10 % FCS passed through the 300K centrifugal filter device after UC scored better in this assay than cells cultivated with 10 % FCS using the 100K centrifugal filter. **(D)** LCLs were cultivated as in panel C but with different % of FCS filtered with 300K centrifugal filters. Results with B cells from two independent donors are shown. RPMI1640 medium supplemented with 2 % FCS after UC and 300K ultrafiltration was used in all experiments throughout the manuscript.

**Supplementary Figure S2**

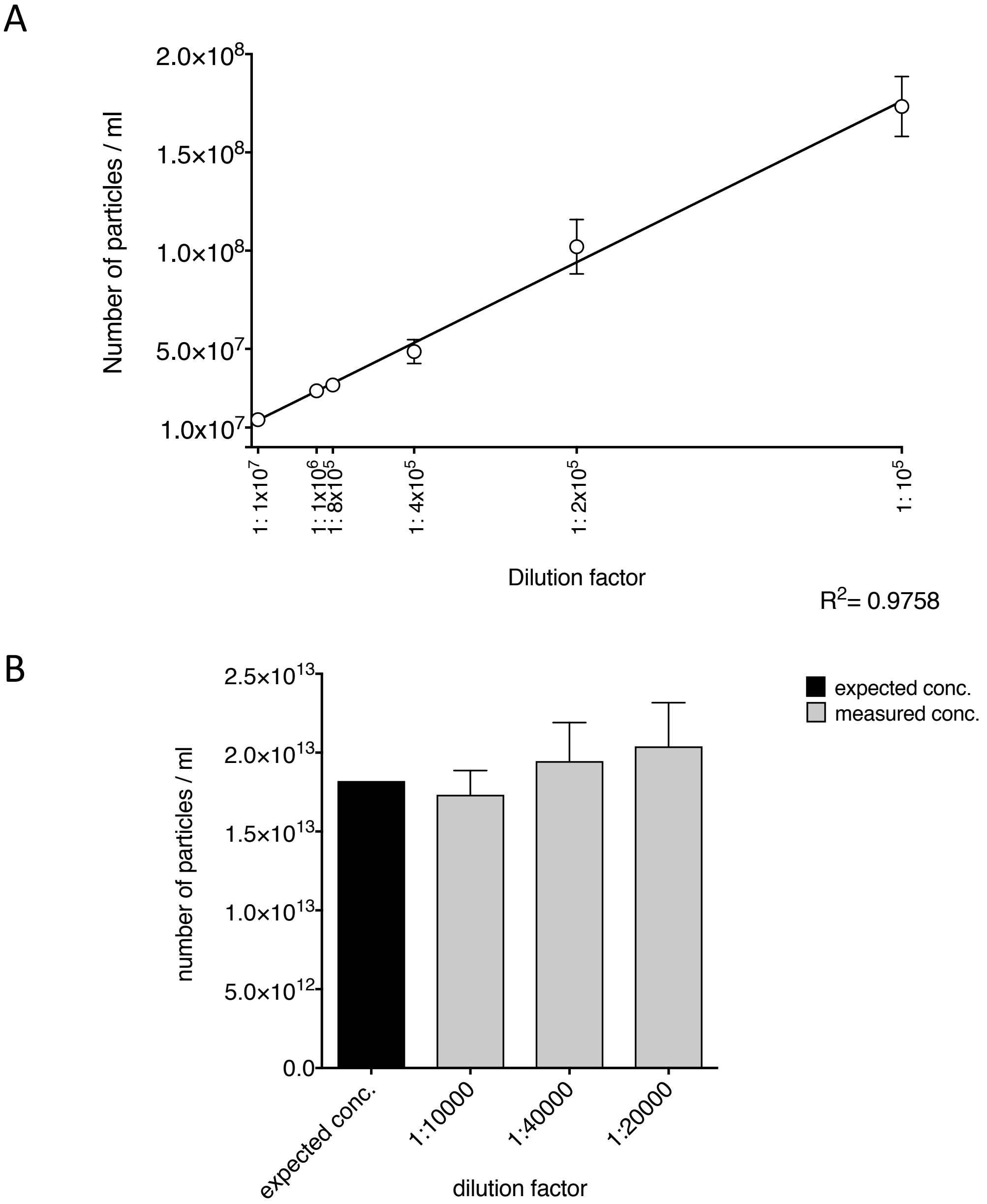


**Validation of the ZetaView PMX110 instrument used for nanoparticle tracking analysis (NTA).**

To assess the accuracy and sensitivity of physical particle measurement with the ZetaView instrument (ParticleMetrix), we used calibration beads of known size (102.7±1.3 nm, Polysciences, Cat. #64010) in the range of EVs. **(A)** Serial dilutions of beads ranging from $1x10^5$ to $1x10^7$ per ml were performed, and each dilution was measured in the ZetaView instrument to confirm the linearity of data acquisition and measurement within this range. **(B)** Accuracy of absolute particle quantification was determined by NTA using dilutions of the calibration beads as in panel A of known initial particle concentration. Mean and standard deviation of three independent replicates are shown.

**Supplementary Figure S3**

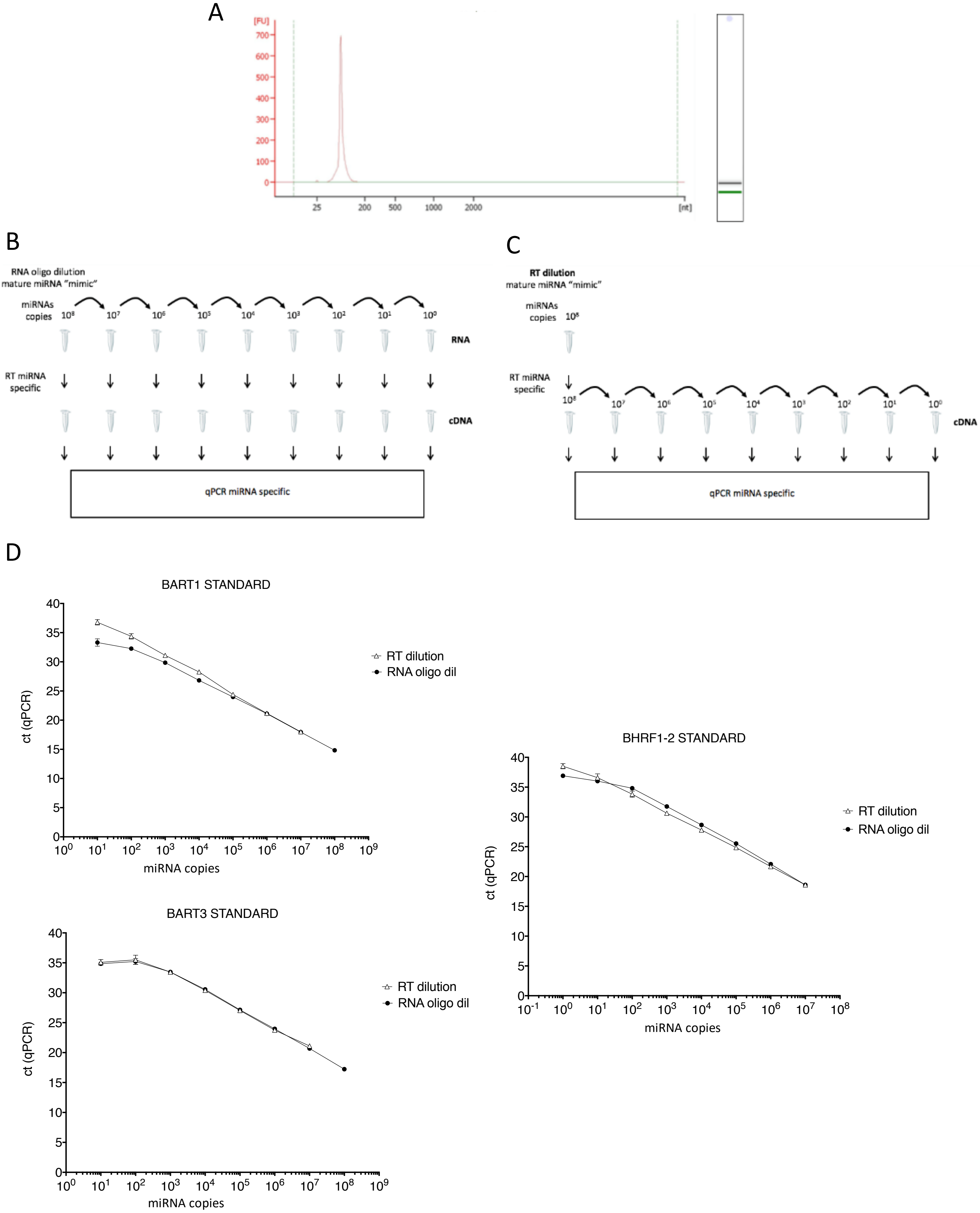

**Absolute quantification of miRNAs.**

For absolute quantification of mature miRNAs, synthetic RNA oligomers ('mimics') corresponding to the mature miRNA sequences of interest were purchased (Metabion). **(A)** The quality of the synthetic RNA molecules was confirmed using an Agilent 2100 Bioanalyzer. Shown is an example of the viral miRNA ebv-miR-BHRF1-2. **(B, C)** Two alternative methods to measure the absolute concentrations of the synthetic miRNAs were compared. **(B)** Single synthetic miRNAs oligomers were diluted first with nuclease-free water to concentrations ranging from $10^0$ to $10^8$ copies per vial followed by reverse transcription (RT) reactions for each vial as described in Materials and Methods. **(C)** Reverse transcription of miRNAs RNA oligomers was performed first, and then the samples were diluted with nuclease-free water to concentrations ranging from $10^0$ to $10^8$ copies. **(D)** After reverse transcription of the miRNAs RNA oligomers as described in panels B or C, quantitative PCRs of each dilution step were performed with three miRNAs of interest: ebv-miR-BART1-5p, ebv-miR-BHRF1-2-3p, and ebv-miR-BART3. The two different approaches of absolute quantification of synthetic miRNAs delivered almost identical results. To mimic the experimental situation, we decided to use the approach shown in panel B in which miRNA samples are diluted prior to reverse transcription. The approach shown in panel C (dilution of a single miRNA sample after reverse transcription) is common, but it does not reflect the situation in samples with only a few miRNA copies in the probes. Mean and standard deviation of three or more independent experiments are shown.

### Supplementary Figure S4

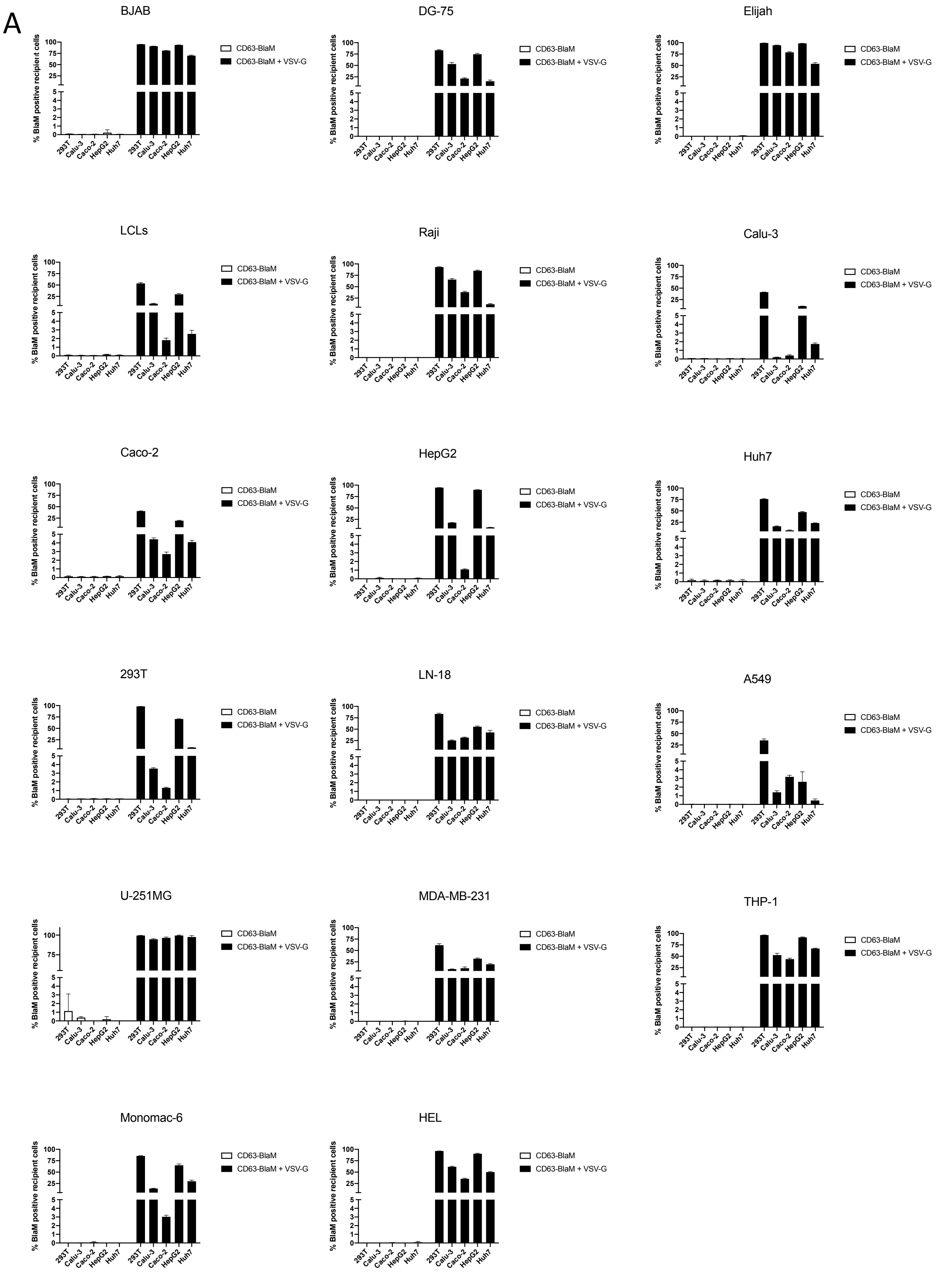

B

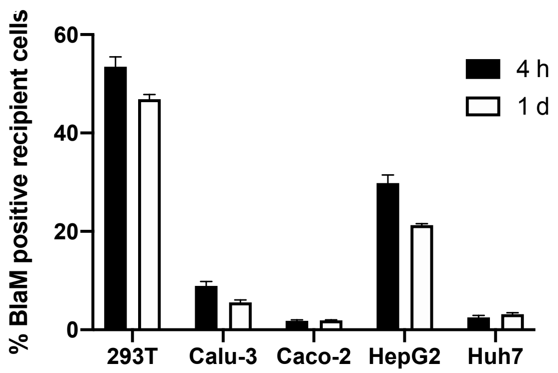


C

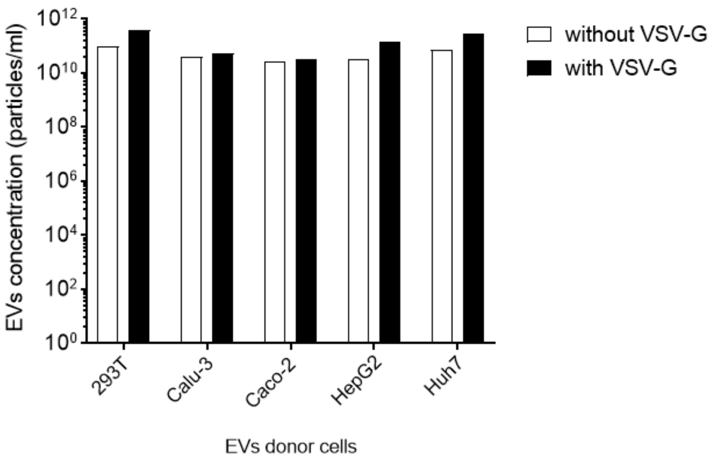


**EVs do not deliver their cargo to recipient cells unless they carry a fusogenic glycoprotein.**

**(A)** 293T, Calu-3, Caco-2, HepG2 and Huh7 were engineered to express CD63-BlaM stably after lentiviral transduction. To boost expression of the CD63-BlaM fusion protein further the cells were transiently transfected with a plasmid expressing the CD63-BlaM fusion protein reporter alone or together with a VSV-G encoding expression plasmid. VSV-G assembled EVs served as positive control. EVs from these cells were purified and $2\times10^5$ cells from 17 different recipient cell lines were incubated for 4 h. The cells were loaded with CCF4-AM substrate, fixed and analyzed by flow cytometry. Means of three replicates are shown. The summary of the data is shown in a heat map in Figure 5E. **(B)** EVs from the 5 different donor cells were generated as in panel A to carry CD63-BlaM and VSV-G and recipient 293T cells were incubated for 4 hours or for one day. Then, the cells were loaded with CCF4-AM substrate, fixed and analyzed by flow cytometry. The mean of three replicates is shown. Differences between cells incubated with EVs for 4 or 24 hours were minor. **(C)** EVs were purified from five different donor cells and physical concentrations of EVs were determined by NTA. 50 µl of each of these preparations was used in Figure 5E.

**Supplementary Figure S5**

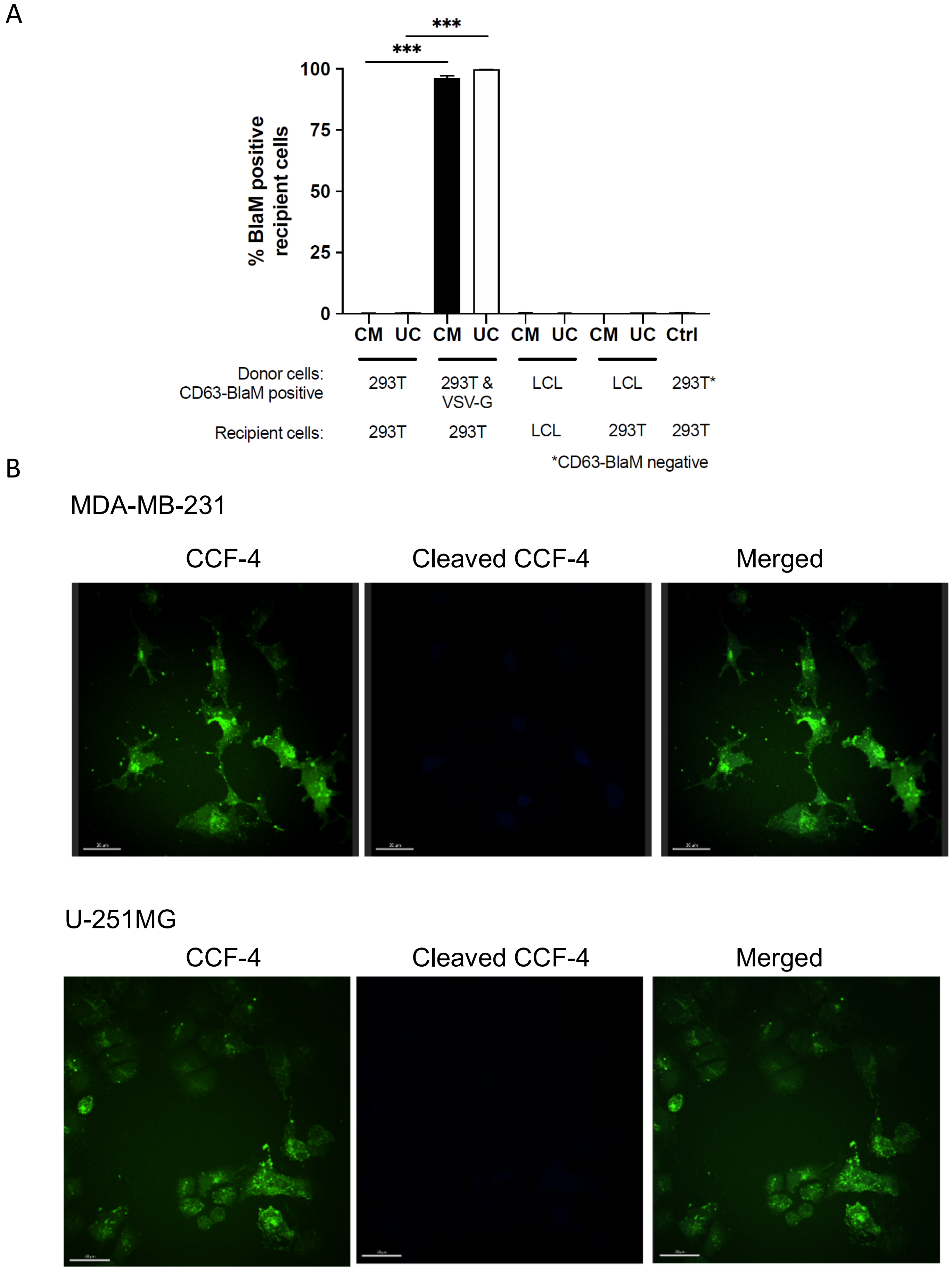

**Details and additional data covering the EV fusion assay.**

(**A**) EBV-positive LCL or 293T cells were engineered to express CD63-BlaM stably after lentiviral transduction. VSV-G was transiently transfected into CD63-BlaM-positive 293T cells as indicated in one case. 500 µl of conditioned medium (CM; see Fig. 2A) or 50 µl resuspended EVs from the UC pellet (UC) were incubated with $2\times10^5$ 293T cells or LCLs as recipients. The negative control (Ctrl) is a sample of 50 µl EVs obtained from an UC pellet with supernatant of non-transduced, parental 293T cells. (**B**) The adherent cell lines U-251MG and MDA-MB-231 were seeded onto glass coverslips (Carl Roth) coated with fibronectin (Advanced Biomatrix). The cells were treated as in Figure 5G but without incubating them with EVs. After 24 hours, medium was replaced with fresh medium and 4 hours later cells were washed three times with PBS and stained with CCF4 overnight. Thereafter, cells were washed three times with PBS, fixed for 10 min with 4 % PFA at room temperature and washed again. The coverslips were mounted with ProLong™ Diamond Antifade Mountant (Thermo Fischer Scientific). The cells were used in parallel as negative controls accompanying panel G of Figure 5. Scale bars is 30 µm.

### Supplementary Figure S6

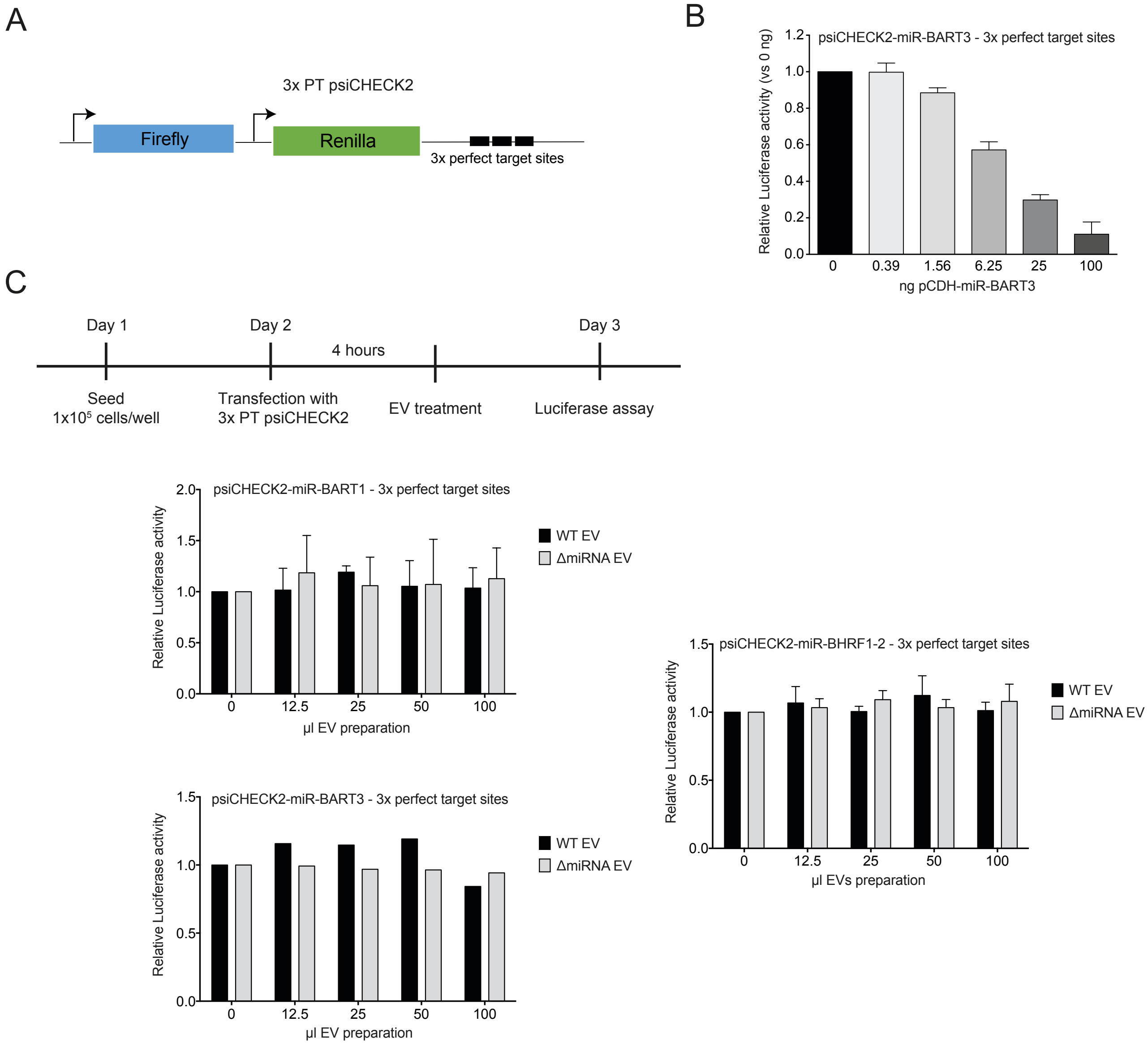


**EVs with EBV miRNAs are not functional in 293T target cells.**

**(A)** The design of the modified dual luciferase reporter plasmid based on psiCHECK2 is shown. It encompasses the internal control firefly luciferase (used for normalization) and the reporter Renilla luciferase with three tandem copies of perfect complementary target sites (3xPT) of the miRNAs of interest inserted in the 3'UTR of the Renilla mRNA. **(B)** 293T cells were transfected with 30 ng of the miRNA reporter plasmid containing 3xPT with increasing amounts of the corresponding miRNA expression vector (pCDH) starting with 390 pg up to 100 ng. At 24 h after transfection, cells were lysed to determine the Renilla and firefly luciferase activities. Mean and SD of three replicates are shown. **(C)** 293T cells were transiently transfected with 30 ng of the 3xPT miRNA reporter plasmid. After 4 h, the cells were incubated with increasing amounts of EVs isolated from the supernatants of LCLs infected with wild-type EBV encoding 44 viral miRNAs (WT EV) or infected with ΔmiRNAs EBV, devoid of all viral miRNAs (ΔmiRNA EV). EVs were prepared from the 'miniUC pellet´ (Fig. 1A) and resuspended in 100 µl, corresponding to approximately $1x10^{11}$ physical particles per ml as measured by NTA. After 24 h incubation, the cells were lysed, and Renilla and firefly luciferase activities were measured. One example of three independent experiments is shown.

**Supplementary Figure S7**

A

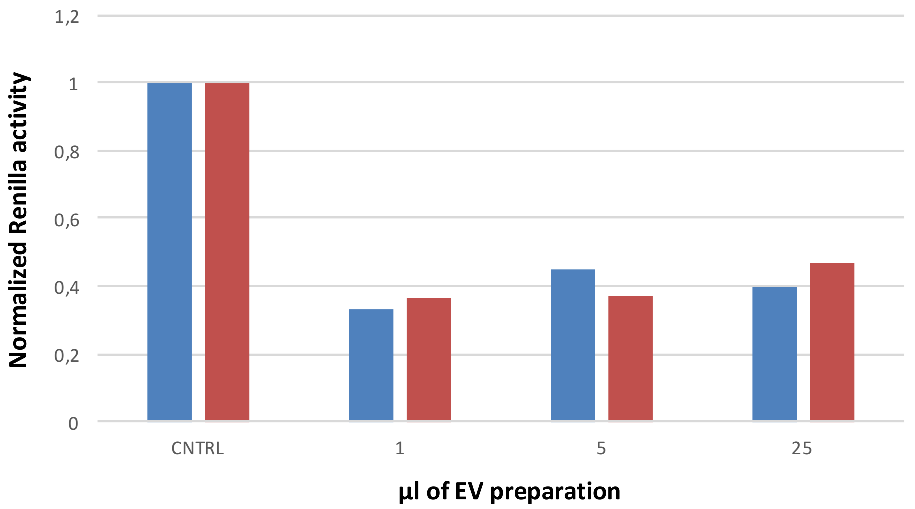


B

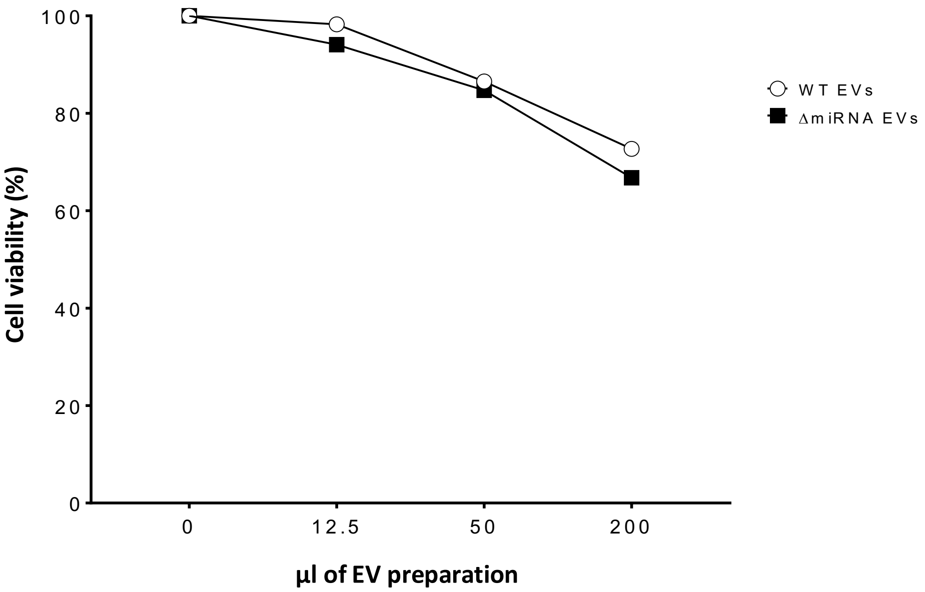


**High doses of LCL-derived EVs are toxic to recipient cells.**

**(A)** 293T cells were seeded in a 96-well plate at an initial density of $2.5\times10^4$ cells/well. After 24 hours, the cells were transiently transfected with 7.5 ng of a miRNA reporter plasmid (3x PT psiCHECK2). Different amounts of EVs were prepared and concentrated from supernatants of WT (blue) or ΔmiRNA EBV (red)-infected B cells were added as indicated on the X-axis 8 h after transfection. 1 µl corresponds to $1\times10^9$ EVs as determined by NTA. After a 24-h incubation, the cells were lysed, and Renilla and firefly luciferase activities were measured. **(B)** 293T cells were seeded in a 24-well plate at an initial density of $2.5\times10^5$cells/well. After 24 hours, the cells were treated with different amounts of EVs isolated from supernatants of WT or ΔmiRNA EBV infected B cells as in panel A. 200 µl contain $2\times10^{10}$ EVs as measured by NTA. After 24 h, cell viability was measured in an MTT assay (Materials and Methods). Cells not treated with EV (0 µl) were set to 100 % for data normalization. Data obtained from one experiment of two independent experiments are shown.
